# SARS-CoV-2 Envelope (E) Protein Interacts with PDZ-Domain-2 of Host Tight Junction Protein ZO1

**DOI:** 10.1101/2020.12.22.422708

**Authors:** Ariel Shepley-McTaggart, Cari A. Sagum, Isabela Oliva, Elizabeth Rybakovsky, Katie DiGuilio, Jingjing Liang, Mark T. Bedford, Joel Cassel, Marius Sudol, James M. Mullin, Ronald N. Harty

**Affiliations:** Department of Pathobiology, School of Veterinary Medicine, University of Pennsylvania, Philadelphia, Pennsylvania, USA; Department of Epigenetics & Molecular Carcinogenesis, M.D. Anderson Cancer Center, University of Texas, Smithville, Texas, USA; The Wistar Cancer Center for Molecular Screening, The Wistar Institute, Philadelphia, PA, USA; Lankenau Institute for Medical Research, Wynnewood, Pennsylvania, USA; Department of Medicine, Icahn School of Medicine at Mount Sinai, New York, New York, USA

**Keywords:** SARS-CoV-2, COVID-19, Envelope (E) protein, ZO-1, PDZ-domain, PDZ binding motif (PBM), tight junction, virus-host interaction, antiviral therapeutic

## Abstract

Newly emerged SARS-CoV-2 is the cause of an ongoing global pandemic leading to severe respiratory disease in humans. SARS-CoV-2 targets epithelial cells in the respiratory tract and lungs, which can lead to amplified chloride secretion and increased leak across epithelial barriers, contributing to severe pneumonia and consolidation of the lungs as seen in many COVID-19 patients. There is an urgent need for a better understanding of the molecular aspects that contribute to SARS-CoV-2-induced pathogenesis and for the development of approaches to mitigate these damaging pathologies. The multifunctional SARS-CoV-2 Envelope (E) protein contributes to virus assembly/egress, and as a membrane protein, also possesses viroporin channel properties that may contribute to epithelial barrier damage, pathogenesis, and disease severity. The extreme C-terminal (ECT) sequence of E also contains a putative PDZ-domain binding motif (PBM), similar to that identified in the E protein of SARS-CoV-1. Here, we screened an array of GST-PDZ domain fusion proteins using either a biotin-labeled WT or mutant ECT peptide from the SARS-CoV-2 E protein. Notably, we identified a singular specific interaction between the WT E peptide and the second PDZ domain of human Zona Occludens-1 (ZO1), one of the key regulators of TJ formation/integrity in all epithelial tissues. We used homogenous time resolve fluorescence (HTRF) as a second complementary approach to further validate this novel modular E-ZO1 interaction. We postulate that SARS-CoV-2 E interacts with ZO1 in infected epithelial cells, and this interaction may contribute, in part, to tight junction damage and epithelial barrier compromise in these cell layers leading to enhanced virus spread and severe respiratory dysfunction that leads to morbidity. Prophylactic/therapeutic intervention targeting this virus-host interaction may effectively reduce airway barrier damage and mitigate virus spread.

## Introduction

Members of the *Coronaviridae* family are enveloped with a positive-sense single-stranded RNA genome and helical nucleocapsid [1]. While symptoms in other mammalian species vary, most coronaviruses cause mild respiratory disease in humans [2–4]. However, a highly pathogenic human coronavirus, severe acute respiratory syndrome coronavirus (SARS-CoV-1), emerged in 2003 to cause acute respiratory disease in afflicted individuals [5, 6]. Moreover, in December 2019, SARS-coronavirus 2 (SARS-CoV-2) emerged as the etiological agent of severe respiratory disease, now called Coronavirus Disease 2019 (COVID-19), first identified in patients in Wuhan, Hubei province, China. This initial outbreak has now become a global pandemic and has afflicted over 71 million people and claimed over 1.6 million lives worldwide as of December 2020. The most common symptoms of COVID-19 patients include fever, malaise, dry cough, and dyspnea with severe cases requiring mechanical ventilation in intensive care unit (ICU) facilities for profound acute hypoxemic respiratory failure [7–10]. Therefore, it is imperative to investigate how this novel SARS-CoV-2 interacts with the host to cause such severe disease pathology.

The Spike (S), Membrane (M), Nucleocapsid (N), and Envelope (E) proteins are the four virion structural proteins that are encoded within the 3′ end of the viral RNA genome. The S protein is responsible for entry and membrane fusion. The M protein is most abundant and gives the virion its shape, while the N protein binds and protects the viral genome as part of the nucleocapsid [2, 11]. The multifunctional E protein plays roles in virion maturation, assembly, and egress, and the E protein of SARS-CoV-1 plays a crucial role in infection as shown by attenuated virulence *in vivo* by a SARS CoV-1 virus lacking the E protein. The extreme C-terminal amino acids of the E protein of SARS-CoV-1 contain an important virulence factor, a PDZ domain binding motif (PBM), whose deletion reduces viral virulence [11–13]. Notably, the PBM of SARS-CoV-1 E protein interacts with the PDZ domain of host protein PALS1, a TJ-associated protein, leading to delayed formation of cellular TJs and disruption of cell polarity in a renal epithelial model [14]. Intriguingly, the extreme C-terminal (ECT) sequence of the E protein of SARS-CoV-2 is similar to that of SARS-CoV-1, suggesting that it may also interact with specific host PDZ-domain bearing proteins via this putative PBM. Indeed, recent studies showed that the SARS-CoV-2 E protein exhibited an increased affinity for PALS1 [9, 15].

We sought to determine whether the ECT of SARS-CoV-2 E protein engages specific host PDZ-domain bearing TJ proteins, which may function to enhance disease progression and severity. To this end, we probed a GST-fused array of approximately 100 mammalian PDZ-domains fixed on solid support with biotin-labeled WT or C-terminal mutant peptides from SARS-CoV-2 E protein. Surprisingly, we identified a single, robust and specific interaction between the WT E peptide and PDZ-domain #2 of human Zona Occludens-1 (ZO1), but not between the C-terminal mutant E peptide and ZO1. ZO1 is a key scaffolding protein that organizes the formation and integrity of TJ complexes via its three PDZ domains that promote multiple protein-protein interactions. Specifically, PDZ-domain #2 of ZO1 is necessary to establish the characteristic continuous band of ZO1 and the TJ barrier proteins, occludin and claudin-2, that is critical for the establishment of normal barrier function across an epithelium [16–18, 31]. We confirmed the E-ZO1 interaction by HTRF once again demonstrating that GST-PDZ domain #2 of ZO1 bound to SARS-CoV-2 E WT peptide, but not with the E mutant peptide, in a concentration dependent manner.

Since severe pneumonia and consolidation of the lungs are often symptoms of COVID-19 [7, 8], it is tempting to speculate that the SARS-CoV-2 E protein may interact with host ZO1 to disrupt or damage TJ complexes and barrier function in human airway epithelial barrier cells as a mechanism to enhance virus spread and disease severity. Further investigations into the interaction between SARS-CoV-2 E protein and ZO1 would improve our understanding of SARS-CoV-2 virus-induced lung morbidity and could be utilized to focus treatment strategies. The putative E/ZO1 interface may prove to be a druggable target, and thus serve to therapeutically reduce SARS-CoV-2 transmission or disease pathology.

## Materials and Methods

### Mammalian PDZ-domain array screen

The PDZ domain array consisted of 90 known PDZ domains and nine 14-3-3-like domains from mammalian proteins expressed in duplicate as purified GST fusion proteins in lettered boxes A-J. SARS-CoV-2 E WT (Biotin-SRVKNLNSSRVP<u>DLLV</u>-COOH) or ECT mutant (Biotin-SRVKNLNSSRVP<u>AAAA</u>-COOH) biotinylated peptides (100ug each) were fluorescently labeled and used to screen the specially prepared C-terminal reading array. Fluorescent spots are indicative of a positive peptide-protein interaction.

### Homogenous Time Resolve Fluorescence (HTRF)

The binding of SARS-CoV-2 WT and ECT mutant E peptides (see above) to purified GST-PDZ domain #2 of ZO1 was assessed using HTRF. Both the protein and the biotinylated peptides were serially diluted 1:2 in assay buffer (25 mM HEPES, pH 7.4, 150 mM NaCl, 5 mM MgCl2, 0.005% Tween-20) and pre-bound for 30 min to either anti-GST-terbium conjugated HTRF donor antibody or streptavidin conjugated to d2 HTRF acceptor (CisBio). Serial dilutions of protein and peptides were then incubated together in a matrix format in a final volume of 10 uL in a white, medium binding, low volume 384-well plate. Following a 1 hr incubation, the HTRF signal was measured using the ClarioStar plate reader (BMG Lab Tech). Data for the WT peptide was fit to a one-site saturation binding model using XlFit (IDBS).

### Expression and Purification of GST-ZO1-PDZ-2 fusion protein

Expression and purification of GST fusion protein from pGEX-ZO1-2/3 plasmid was performed as described previously [19, 20]. Briefly, the plasmids were transformed into *E. coli* BL21(DE3) cells and single colonies were cultured in 10ml of LB media overnight with shaking at 37°C. The overnight culture was added into 100ml of fresh LB broth and grown at 37°C for one hour with shaking. GST alone or GST-PDZ domain fusion proteins were induced with isopropyl-β-d-thiogalactopyranoside (IPTG) (0.1 mM) for 4 h at 30°C. Bacterial cultures were centrifuged at 5,000 rpm for 10 min at 4°C, and lysates were extracted by using B-PER bacterial protein extract reagent according to the protocol supplied by the manufacturer (Pierce). Fusion proteins were purified with glutathione-Sepharose 4B and eluted with elution buffer (100 mM Tris-Cl [pH 8.0], 120 mM NaCl, 30 mM reduced glutathione). Purified proteins were analyzed on SDS-PAGE gels and stained with Coomassie blue and quantified.

## Results

### Comparison of C-terminal sequences of SARS-CoV-1 and SARS-CoV-2 E proteins

The SARS-CoV E protein is a small membrane protein that has multiple functions in infected cells and is incorporated into mature virions [21–24]. The E protein can be divided into 3 major regions including N-terminal, trans-membrane, and C-terminal domains (Fig. 1). In addition, the extreme C-terminal amino acids of SARS-CoV-1 E (DLLV) comprise a validated PDZ-domain binding motif (Fig. 1, red). Since the DLLV core motif is perfectly conserved in the SARS-CoV-2 E protein (Wuhan-Hu-1 strain), and the immediately adjacent amino acids, although not identical, are highly conserved with those of SARS-CoV-1 E (Fig. 1), the likelihood that SARS-CoV-2 E protein can also interact with select PDZ-domains of host proteins is high [14, 24–26].

**Fig. 1.**
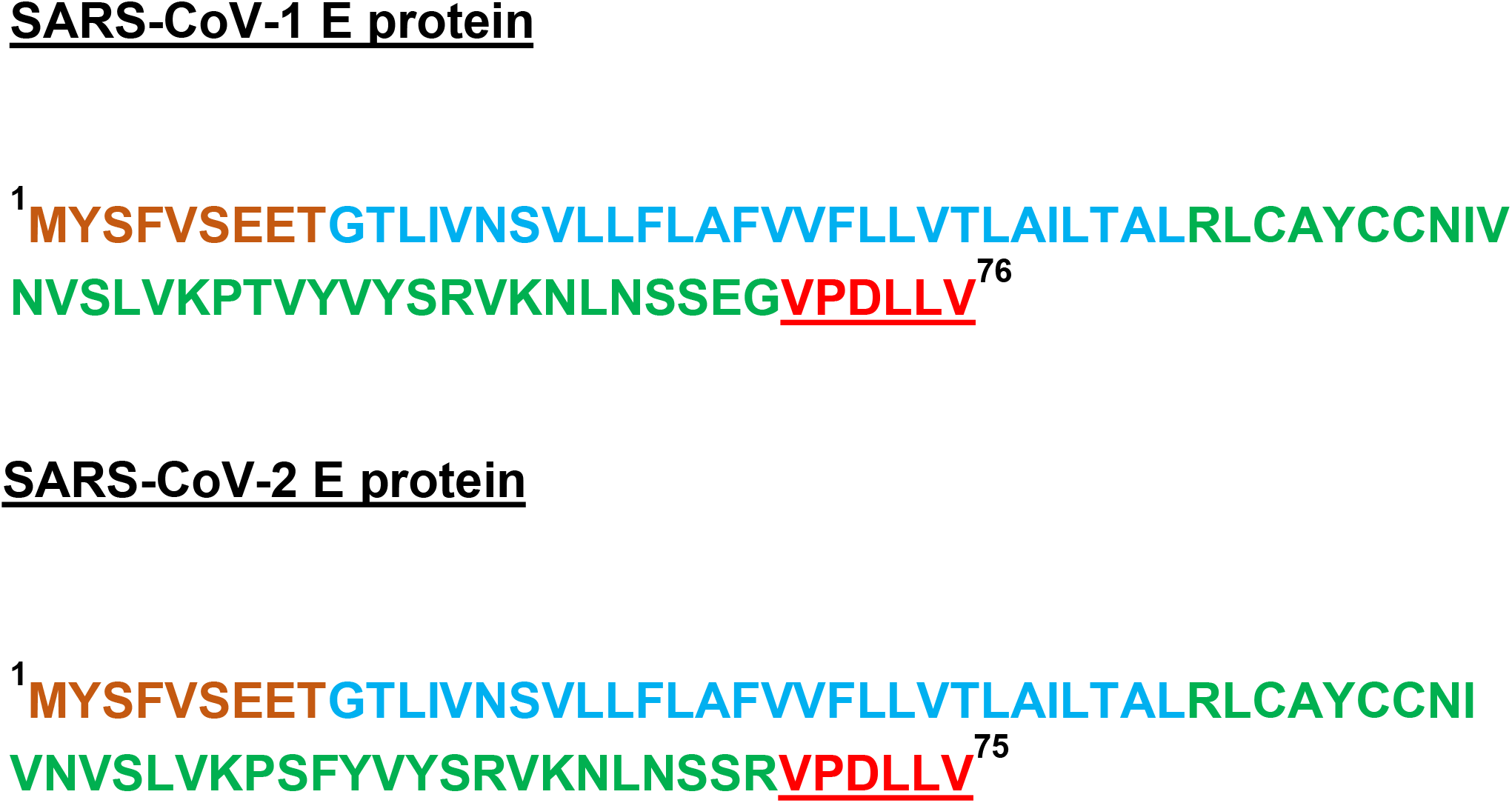
Amino acid sequence of full-length SARS-CoV-1 and SARS-CoV-2 E proteins. The N-terminal (brown), trans-membrane (blue), and C-terminal (green) regions are highlighted. The conserved C-terminal DLLV sequences and putative PBM motifs are shown in red.

### Identification of ZO1 as a PDZ-domain interactor with the E protein of SARS-CoV-2

Here, we sought to determine whether the putative PBM present within the SARS-CoV-2 E protein could interact with a wide-array of host PDZ-domain containing proteins. We used fluorescently-labeled, biotinylated peptides containing WT or mutated C-terminal sequences from the SARS-CoV-2 E protein to screen an array composed of 90 PDZ-domains and 9 14-3-3-like domains derived from mammalian proteins to detect novel host interactors (Fig. 2, top). Surprisingly, we identified a singular specific interaction between the SARS-CoV-2 WT E peptide and the second PDZ-domain of human ZO1 (Fig. 2, bottom left panel, red oval, position J6). As a control, the SARS-CoV-2 mutant E peptide did not interact with any of the PDZ- or 14-3-3-domains present on the array (Fig. 2, bottom middle panel). All GST fusion proteins were present on the array as indicated by the use of anti-GST antiserum (Fig. 2, bottom right panel). To our knowledge, these results are the first to identify PDZ-domain #2 of human ZO1 as an interactor with the C-terminal sequences of SARS-CoV-2 E protein.

**Fig. 2.**
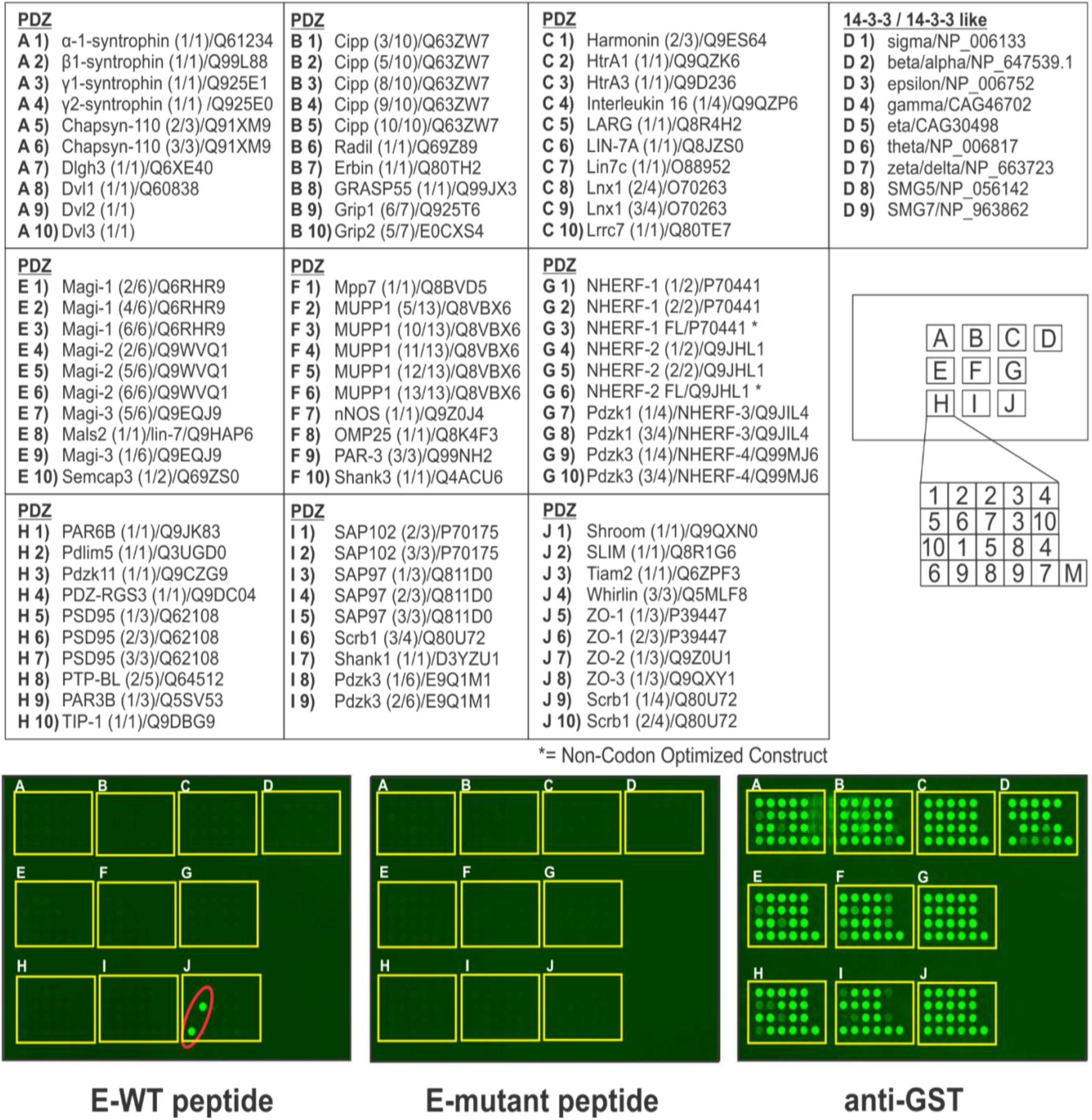
Screening array of GST-PDZ domain fusion proteins. The indicated GST-PDZ and GST-14-3-3 domain fusion proteins per lettered box were arrayed in duplicate as shown. The bottom right sample (M) in each box represents GST alone as a negative control. The array was screened with biotinylated E-WT or E-mutant peptides of SARS-CoV-2 E protein. Representative data for E-WT peptide (Biotin-SRVKNLNSSRVPDLLV) (100μg), and E-mutant peptide (Biotin-SRVKNLNSSRVPAAAA) (100μg) are shown (bottom panels). The E-mutant peptide did not interact with any GST-PDZ or GST-14-3-3 domain fusion proteins, whereas the E-WT peptide interacted strongly and solely with GST-PDZ domain #2 from human ZO1 in position 6 in box J (red oval). A positive control for expression of all GST fusion proteins is shown (anti-GST).

### Use of Homogenous Time Resolve Fluorescence (HTRF) to confirm the E-ZO1 interaction

We sought to use a second, complementary approach to validate the E-ZO1 interaction identified in our screening array. Toward this end, we used an HTRF assay to assess the binding of SARS-CoV-2 WT and ECT mutant E peptides to purified GST-ZO1 PDZ domain #2 (Fig. 3). Serial dilutions of GST-ZO1 PDZ-domain protein and WT or mutant E peptides were incubated together for 1 hour, and the HTRF signal was measured and plotted using XIFit. Indeed, we observed a clear concentration dependent binding of GST-ZO1 PDZ domain #2 to WT (Fig. 3, left panel), but not the ECT mutant (Fig. 3, right panel), E peptide over a range of peptide concentrations. The Kd of the interaction between the SARS-CoV-2 WT E peptide and a 3nM concentration of GST-ZO1 PDZ domain #2 was 29nM (Fig. 3, bottom panel), which is consistent with that calculated for PDZ domains and their cognate peptides. These findings confirm those described above indicating that the C-terminal sequences of SARS-CoV-2 E protein interact robustly with PDZ-domain #2 of human ZO1 protein.

**Fig. 3.**
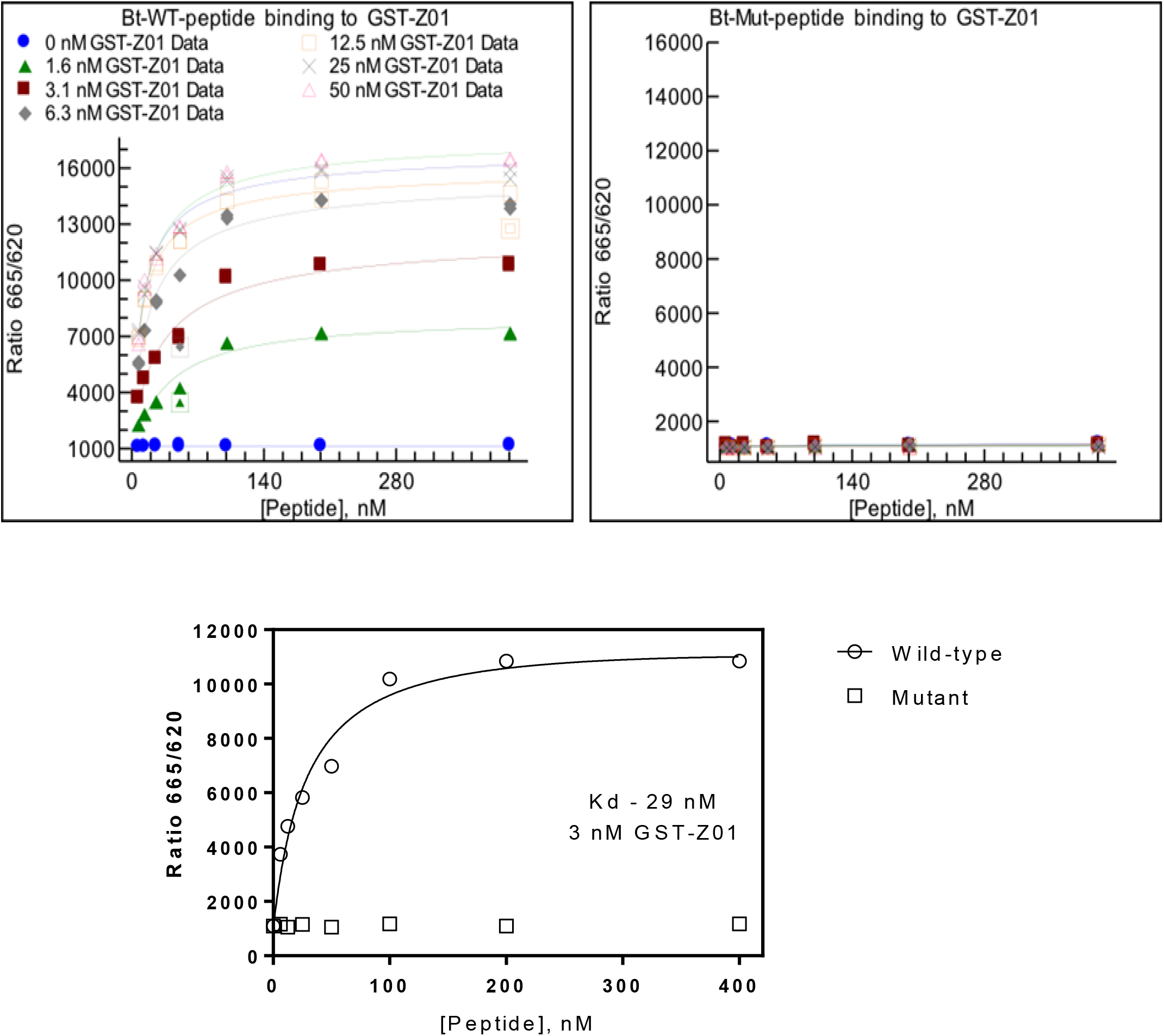
Homogenous Time Resolve Fluorescence (HTRF). The concentration dependent binding properties of SARS-CoV-2 E-WT (Bt-WT peptide, left panel) and E-mutant (Bt-Mut peptide, right panel) peptides with purified GST-PDZ domain #2 of ZO1 are shown. Concentrations of the GST-PDZ domain fusion protein ranging from 0-50 nM (indicated by the various symbols) were incubated with the indicated concentrations of E-WT (left) or E-mutant (right) peptides (x-axis), and the HTRF signal was measured. The E-WT peptide did not bind to GST alone (left panel, blue dots); however, clear concentration dependent binding of the E-WT peptide to GST-PDZ domain 2 of ZO1 was observed. The E-mutant peptide did not bind to any concentration of GST-PDZ domain 2 of ZO1 tested (right panel). A Kd value of 29nM was calculated for E-WT peptide binding to 3nM concentration of GST-ZO1 PDZ domain #2 (bottom panel).

## Discussion

The coronavirus E protein has many functions during infection, including the assembly, budding, and intracellular trafficking of infectious virions from the ER-Golgi complex. Modeling of the SARS-CoV-2 E protein suggests that E can form a broadly cation selective ion channel with dynamic open and closed states, and therefore may act as a viroporin similar to the SARS-CoV-1 E protein [21, 28]. Although the genome of SARS-CoV-2 only shares about 80% identity with that of SARS-CoV-1, the ECT of the SARS-CoV-2 E protein, including the DLLV core motif and adjacent amino acids, are highly conserved with those of SARS-CoV-1 E protein [21–23, 25, 27] (Fig. 1). Sequence alignment of the SARS-CoV-1 and SARS-CoV-2 E proteins (Fig 1) highlight the sequence similarities and the putative PBM located at the ECT in both proteins.

We utilized a screening array to identify possible host PDZ-domain containing proteins that may interact with the putative PBM of the SARS-CoV-2 E protein. Excitingly, we identified a single positive hit, PDZ domain #2 of TJ scaffolding protein ZO1 (Fig 2). We confirmed this novel interaction between the SARS-CoV-2 E WT peptide and purified GST-PDZ domain #2 of ZO1 using HTRF (Fig 3). To our knowledge, these data are the first to identify PDZ-domain #2 of ZO1 as an interactor with the ECT of the SARS-CoV-2 E protein.

There is precedent for coronavirus E proteins interacting with host PDZ domains. Indeed, the SARS-CoV-1 E protein was shown to interact with the PDZ domain of TJ protein PALS1, resulting in mislocalization of PALS1 and delayed formation of TJs in a renal model. In a subsequent study, the C-terminal sequences of SARS-CoV-2 E protein were also predicted to bind to the PDZ domain of PALS1 [14, 15]. We did not detect an E-PALS1 interaction, since the PDZ domain of PALS1 was not included in the screening array (Fig 2). In addition, PDZ-domain #2 of ZO2 and ZO3 proteins were also missing from our screening array. Importantly, the second PDZ domains of ZO1, ZO2, and ZO3 are fairly-well conserved in their structures, and they homo- and/or hetero-dimerize to form functional multi-component complexes of ZO proteins in tight junctions. Therefore, we speculate that the second PDZ domains of ZO2 and ZO3 would also interact positively with the ECT of the SARS-CoV-2 E protein.

Several pathogens, including respiratory viruses, may cause breakdown of cellular barriers (as well as cell polarity and tissue-specific unidirectional transport processes) in the lung by a mechanism involving an interaction between the virus and lung epithelial cell TJs to induce leak. These pathogens can interact with cellular proteins comprising TJ complexes, thereby causing their disruption and subsequently enhancing systemic virus spread across epithelial and endothelial barriers [29, 30] (Fig. 4). There are several independent mechanisms by which an E-ZO1 interaction may disrupt TJs, barrier integrity, physiologically vital transcellular transport processes, and possibly cell polarity. For example, SARS-CoV-2 E protein may compete with other proteins of the TJ complex (*e.g*. ZO2, ZO3, or ZO1 itself [homodimerizing]) for binding to PDZ domain #2 of ZO1, leading to the inability of claudins to organize into actual barrier-forming strands [18]. Alternatively, an E-ZO1 interaction could potentially alter ZO1 binding to actin filaments of the actin-myosin ring, thereby altering barrier regulation through myosin light chain kinase signaling. In addition, an E-ZO1 interaction at the TJ complex could result in a decreased affinity for ZO1 binding to ZONAB (ZO1-associated nucleic acid binding protein), dislocating ZONAB from the TJ complex and increasing ZONAB translocation to the nucleus, leading to an increase of epithelial mesenchymal transition (EMT) and a concomitant decrease in epithelial barrier function (as well as unidirectional chloride secretion) through general epithelial dedifferentiation [31–33].

**Fig. 4.**
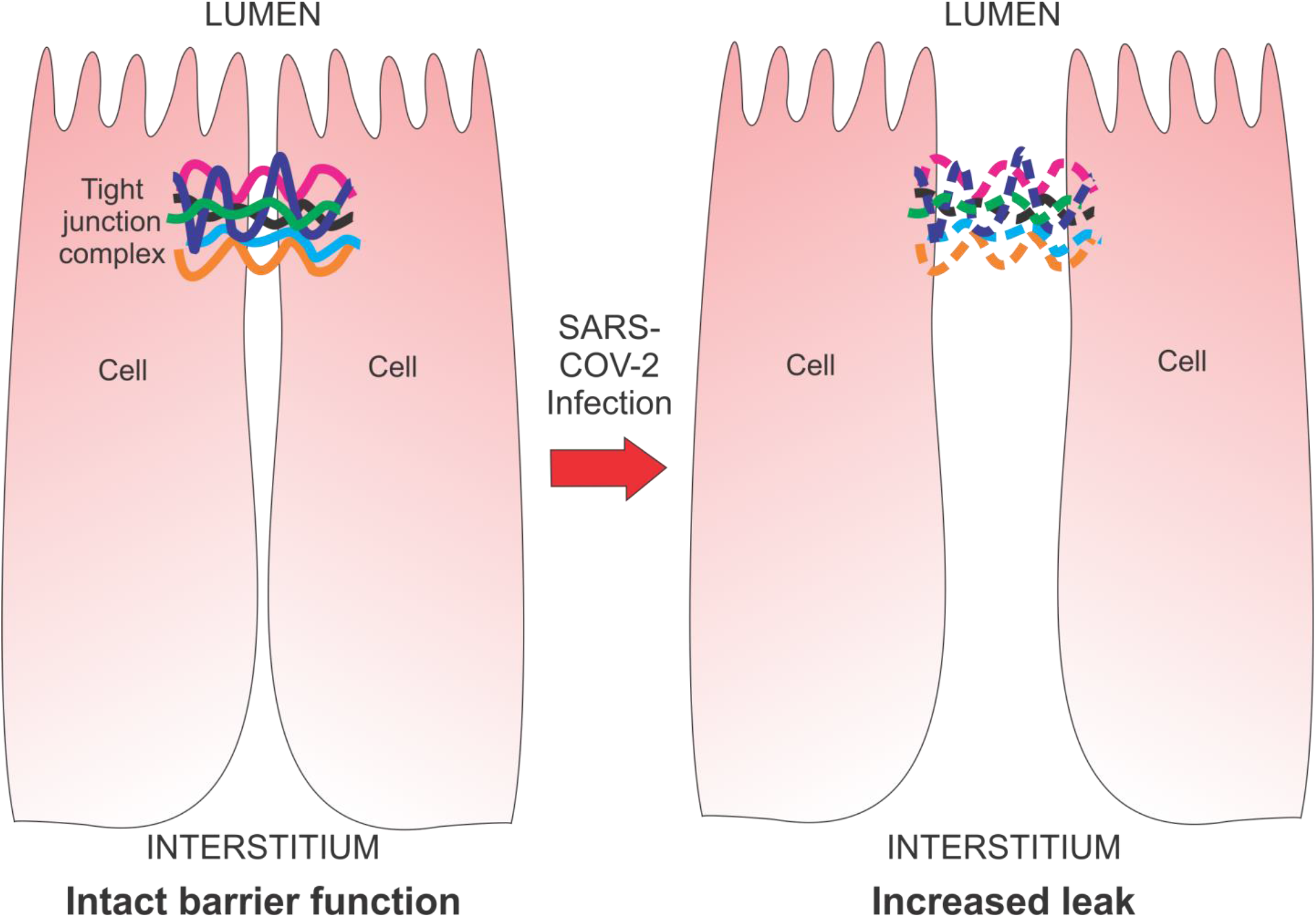
Model diagram of potential damage to tight junction integrity and epithelia following infection by SARS-CoV-2. **Left** – intact epithelial TJ complex and barrier function from lumen to interstitium. Cells are close together with solid colored lines representing TJ proteins ZO1, ZO2, ZO3, and PALS1. **Right** – compromised epithelial TJ complex (dotted colored lines) following SARS-CoV-2 infection and expression of E protein resulting in barrier disruption and increased paracellular transepithelial leak.

By disrupting TJ integrity, the E-ZO1 interaction would result in increased paracellular transepithelial leak as well as altered unidirectional transcellular salt and water transport, contributing to accumulation of water in the lungs of COVID19 patients. In fact, a recent study shows that SARS-CoV-2 infection disrupts ZO1 localization, and causes barrier dysfunction as demonstrated by a decrease in transepithelial electrical resistance (TEER) [34]. The authors speculate that this decrease in barrier function in infected cells may be due to cytokine release in a secondary inflammatory response. It remains to be determined whether an E-ZO1 interaction represents a more primary mechanism that contributes to this observed decrease in barrier function and altered localization of ZO1 in infected cells as described above.

Ongoing studies will determine whether full length SARS-CoV-2 E interacts with endogenous ZO1 in virus infected cells, in an effort to establish the biological significance of this virus-host interaction. In addition, experiments are underway with lentivirus particles engineered to express the SARS-CoV-2 WT and mutant E proteins alone in human lung airway cells (Calu-3 and 16HBE) in order to investigate the E/ZO1 interaction in isolation and its potential effect on TJ integrity via TEER analysis and transepithelial diffusion of paracellular probe molecules. Validation of the E/ZO1 interaction in virus infected cells will be important for future development of small molecule compounds to therapeutically lessen lung morbidity and disease symptoms in severe COVID19 patients. Targeting the E/ZO1 interface for treatment may effectively reduce airway barrier cell damage and diminish the morbidity and mortality associated with this major health threat.

## Acknowledgements

The authors thank D. Argento for illustrations and graphics. Funding was provided in part from a University of Pennsylvania School of Veterinary Medicine COVID-19 Pilot Award, a University of Pennsylvania School of Medicine Institute for Translational Medicine and Therapeutics Pilot Award, a Fast Grant from the Emergent Ventures Program at the Mercatus Center at George Mason University, and NIH NIAID awards AI138052 + AI139392 to R.N.H, and a Fast Grant from the Emergent Ventures Program at the Mercatus Center at George Mason University to J.M.M. A.S-M. was supported in part by T32-AI070077 award from NIH/NIAID. Probing of arrayed methyl reader domains was made possible via the UT MDACC Protein Array & Analysis Core (PAAC) CPRIT Grant RP180804 (Directed by M.T.B). The funders had no role in study design, data collection and analysis, decision to publish, or preparation of the manuscript.

